# Memory Guides the Comprehension of Event Changes for Older and Younger Adults

**DOI:** 10.1101/201939

**Authors:** Christopher N. Wahlheim, Jeffrey M. Zacks

## Abstract

Two experiments examined adult age differences in the use of memory to comprehend changes in everyday activities. Participants viewed movies depicting an actor performing activities on two fictive days in her life. Some activities were repeated across days, other activities were repeated with a changed feature (e.g., waking up to an *alarm clock* or a *phone alarm*), and a final set of activities was performed on Day 2 only. After a one-week delay, participants completed a cued recall test for the activities of Day 2. Unsurprisingly, exact repetition boosted final recall. More surprising, features that changed from Day 1 to Day 2 were remembered approximately as well as features that were only presented on Day 2—showing an absence of proactive interference and in some cases proactive *facilitation*. Proactive facilitation was strongly related to participants’ ability to detect and recollect the changes. Younger adults detected and recollected more changes than older adults, which in part explained older adults’ differential deficit in memory for changed activity features. We propose that this pattern may reflect observers’ use of episodic memory to make predictions during the experience of a new activity, and that when predictions fail, this triggers processing that benefits subsequent episodic memory. Disruption of this chain of processing could play a role in age-related episodic memory deficits.

---

Many of the activities people experience are not brand new, and not completely novel, but are near repetitions—variations on a theme. For example, suppose you had a friend who always ordered a cheeseburger for dinner, but then discovered a family history of heart disease. When you next have dinner with this friend, you might predict that she would place her regular order, but then experience a prediction error if she ordered a salad. Registering this change, consciously or not, would help you to better predict her behavior the next time you eat together. The processing associated with registering the change might also affect your encoding of this particular episode. In this paper, we develop a theoretical framework that proposes a mechanism by which people use memory for recent past events to guide their comprehension and encoding of the present, with consequences for subsequent memory. This framework is based on previous studies of memory for change and previous research on event comprehension. It leads to predictions about how memory for changes in events is affected by aging, which we tested in two experiments. Before describing the framework and the present experiments, we describe their origins in studies of event perception, episodic memory, and cognitive aging.

## Event Perception and Memory

People perceive naturalistic ongoing activities as discrete events. Theories of event perception hold that everyday activities are represented hierarchically (e.g., Dickman, 1963; Brewer & Dupree, 1983; Zacks & Tversky, 2001), such that smaller units of activity are nested within larger units of activity. For example, the event described above (eating dinner) is comprised of smaller events such as ordering food and eating it, which in turn might be broken down further into events such as picking up a fork, cutting a burger, and taking a drink from a glass of water. Understanding structure in events allows individuals to establish better organized representations of what is happening in their current environment. People spontaneously segment activities during event perception (e.g., Zacks et al., 2001), and this is related to their subsequent memory: Features from event boundaries are often remembered better than event middles (Newtson & Engquist, 1976; Schwan, Garsoffky, & Hesse, 2000; Swallow, Zacks, & Abrams, 2009), individual differences in event segmentation predict individual differences in event memory (Sargent et al., 2013; Zacks, Speer, Vettel, & Jacoby, 2006; Flores, Bailey, Eisenberg, & Zacks, 2017), and interventions to facilitate effective segmentation causally improve memory (Boltz, 1992; Flores et al., 2017; Gold, Zacks, & Flores, 2016; Schwan et al., 2000).

Event Segmentation Theory (EST; Zacks, Speer, Swallow, Braver, & Reynolds, 2007) gives an account of how ongoing activity is segmented into meaningful chunks, with implications for episodic memory formation. According to EST, observers form working memory representations of an ongoing activity, called *event models*, that represent “what is happening now.” These models serve to guide the comprehension of incoming perceptual information. An effective current event model allows one to make predictions about what will happen in the near future. When there is a spike in prediction error, the current event model is updated. This process of memory updating is experienced by the observer as a boundary between events. Of particular relevance to the present study is the notion that prediction plays a central role in establishing memory representations of discrete units of activity.

The role of prediction in the encoding of event representations has also been discussed in the episodic memory literature. For example, Glenberg (1997) has argued that episodic memories allow individuals to predict physical interactions with their three-dimensional worlds. Related to this, Hintzman (2011) has suggested that involuntary recollections of past events cue potential spatio-temporal regularities and that such remindings can facilitate predicting future events.

Finally, work on *episodic future thought* suggests that mental representations underlying episodic memory also enable thinking predictively about future events (Schacter, Addis, Hassabis, Martin, Spreng, & Szpunar, 2012). In this vein, a recent study of the neurophysiology of event segmentation and memory indicated that prior memories guide anticipatory reinstatement of neural activity in regions associated with segmentation at the level of event models (e.g., Baldassano, Chen, Zadbood, Pillow, Hasson, & Norman, 2017). These findings suggest that episodic memory representations could facilitate detection of changes from one instance of an event type to the next; however, no previous studies have investigated this.

## Detecting and Recollecting Episodic Changes

Episodic changes and their attendant effects on memory have been investigated extensively in the episodic memory literature, in the context of interference effects. It is often the case that episodic changes impair memory when different features are associated with a common cue. For example, in our restaurant example the two meals are different features associated with a common cue, the restaurant. Looking back, you might have difficulty remembering which meal your friend ate at the restaurant on the second dinner because you could experience *proactive interference* from the memory of the first dinner. Indeed, the deleterious effects of such response competition have been demonstrated across many paradigms using a variety of stimulus materials (for a review, see Anderson & Neeley, 1996). However, there are some situations in which multiple associations can facilitate remembering. In a classic example, Barnes and Underwood (1959) found *retroactive facilitation* in paired associate learning when cues were presented with associated responses in separate lists (A-B, A-B’). Consistent with this, Robbins and Bray (1974) found retroactive facilitation in an A-B, A-D paradigm in which participants were told about the relationship between pairs in each list (also see, Bruce & Weaver, 1973). Findings such as these have been taken to suggest that episodic changes can lead to enhanced memory when individuals detect and encode those changes.

Following this proposal, Jacoby, Wahlheim and colleagues (e.g., Jacoby, Wahlheim, & Kelley, 2015; Wahlheim & Jacoby, 2013) proposed the memory-for-change framework to explain when episodic changes should enhance or impair memory. A key component of this framework is the recursive reminding hypothesis (Hintzman, 2011), which states that perceiving a current event can trigger involuntary recollection of past events, and the cognitive operation of reminding can be encoded as part of a configural representation that also includes the constituent events of the reminding (also see, Hintzman, Summers, & Block, 1975; Tzeng & Cotton, 1980; Winograd & Soloway, 1985). This representation is recursive because memory for a later event becomes embedded within a trace that includes reminding of earlier events. This attribute can facilitate order memory because more recent events embed earlier events, but not vice versa.

The empirical findings taken as initial support for the recursive reminding hypothesis were those showing effects of repetitions and item associations on spacing judgments (Hintzman & Block, 1973; Hintzman et al., 1975), effects of spaced repetitions on frequency judgments (Hintzman, 2004) and effects of item associations on relative order judgments (Hintzman, 2010). Jacoby, Wahlheim, and colleagues (e.g., Jacoby et al., 2015; Wahlheim & Jacoby, 2013) extended this hypothesis by proposing a dual-process model that integrates controlled and automatic influences of memory with the mechanics proposed by the recursive remindings hypothesis. According to their model, the retrieval processes involved in detecting a changed event brings the earlier event into the same context as the current event that contains the changed features. In doing so, change detection allows both events to be encoded into a configural trace that also includes memory for the retrieval event (i.e., reminding). At test, recollecting change provides access to the temporal order of events by allowing individuals to remember that the recent event reminded them of the earlier event. In contrast, failing to recollect earlier detected changes results in impaired memory for the more recent event because the accessibility of competing responses is heightened by the retrieval that precedes change detection, and this accessibility is unopposed when retrieval at test is not based on change recollection.

A clear example of the predicted effects of the memory-for-change framework can be seen in a paired-associate learning experiment from Jacoby et al. (2015, Experiment 3). Participants studied two lists of words that contained pairs that changed within List 2, and pairs that changed between presentations in List 1 and in List 2. Participants were told either to look for all the possible changes in the experiment, or to look only for changes within List 2. The critical question was how participants performed on a final test of memory for the List 2 pairs when items changed from List 1 to List 2. Those who had looked for all possible changes performed much better on these pairs, and were better able to recollect that the item had changed across lists. These results showed direct evidence that making contact between changed events can enhance memory in the manner predicted by the memory-for-change account.

Several other studies have shown similar effects of episodic changes using various materials. These effects have been shown in order memory for categorially related pairs of words (Jacoby & Wahlheim, 2013), list discrimination of word pairs (Jacoby, Wahlheim, & Yonelinas, 2013), memory for positions on controversial issues held by fictional political candidates (Putnam, Wahlheim, & Jacoby, 2014), and misinformation about a fictional crime (Putnam, Sungkhasettee, & Roediger, 2017). The latter two results are important for the present study because they suggest that effects of change are likely to generalize to more naturalistic contexts. The most naturalistic materials were used in the study by Putnam et al. (2017), who found that memory for details from a slideshow depicting a fictional crime that were changed in a subsequently presented narrative were better remembered when those changes were recollected at test. This study has high ecological validity and the misinformation paradigm has both theoretical and practical significance. However, for revealing the mechanisms by which change detection affects memory the fact that it tests retroactive effects of memory is a limitation. In this paradigm, memory enhancement resulting from change detection could be the result of retrieval practice rather than change detection.

## Adult Age Differences in Event Perception and Memory for Change

One process that may have profound effects on the ability to use memory for predicting new events is healthy aging. Older adults have well established deficits in episodic memory tasks, especially those that require self-initiated reinstatement of context from prior events (for reviews, see Balota, Dolan, & Duchek, 2000; Zacks, Hasher, & Li, 2000). Research has shown that this deficit extends to older adults' memory for naturalistic activities resulting from poorer encoding of event structure. For example, Zacks, Speer, Vettel, and Jacoby (2006) found that older adults segmented ongoing activities less consistently than younger adults, and this predicted older adults' poorer memory for event details. In addition, Kurby and Zacks (2011) found that older adults' segmentation was less hierarchically organized than younger adults, older adults had worse memory than younger adults, and age differences in event segmentation sometimes predicted these memory differences. Given that the ability to segment events depends on one's sensitivity to changes, these findings suggest that older adults may be less sensitive to changes in ongoing activities. This possibility is consistent with the finding that older adults are less sensitive to changes in context (Balota, Duchek, & Paulin, 1989). Consequently, older adults may form less coherent event representations, which could also impair their ability to detect when event features change from one episode to the next.

In addition to this age-related deficit in detecting ongoing changes, older adults are impaired in their detection of changes that occur between separate episodes. Important for the present study, this deficit has downstream consequences for change recollection and recall under conditions of proactive interference. A clear example of this can be seen in a paired associate learning experiment from Wahlheim (2014; Experiment 3). In this experiment, participants studied two lists of word pairs and were instructed to indicate which pairs changed from List 1 to List 2. Older adults were given extra study time in List 2, which allowed them to detect as many changes as younger adults. Despite this, older adults still recollected fewer changes, and this partly explained why older adults showed proactive interference in their recall, but younger adults did not. Older adults' greater susceptibility to interference shown in that study is similar to results shown in other interference paradigms (e.g., Hasher & Zacks, 1988; Jacoby, Debner, & Hay, 2001; Healey, Hasher, & Campbell, 2013). However, these results from Wahlheim (2014) showed that in situations involving episodic change, this deficit can be partly explained by differences in the ability to detect and recollect changes. When considered in the context of event perception, these findings suggest that older adults' impaired ability to form coherent episodic representations of naturalistic activities could create a deficit in episodic change detection with similar downstream consequences for remembering changes and associated activity features.

## Change Comprehension Theory

Bringing together the role of prediction in event comprehension proposed by EST (Zacks et al., 2007) with the role of change processing in episodic memory proposed by the memory-for-change framework (Jacoby et al., 2015; Wahlheim & Jacoby, 2013) leads to a novel proposal for the mechanism by which memories guide ongoing comprehension, and at the same time determine new episodic memory encoding of event changes. We refer to the account of this mechanism as Change Comprehension Theory (CCT). In this section, we describe CCT and explain its implications for the effects of healthy aging on memory for change described above.

**Figure 1.**
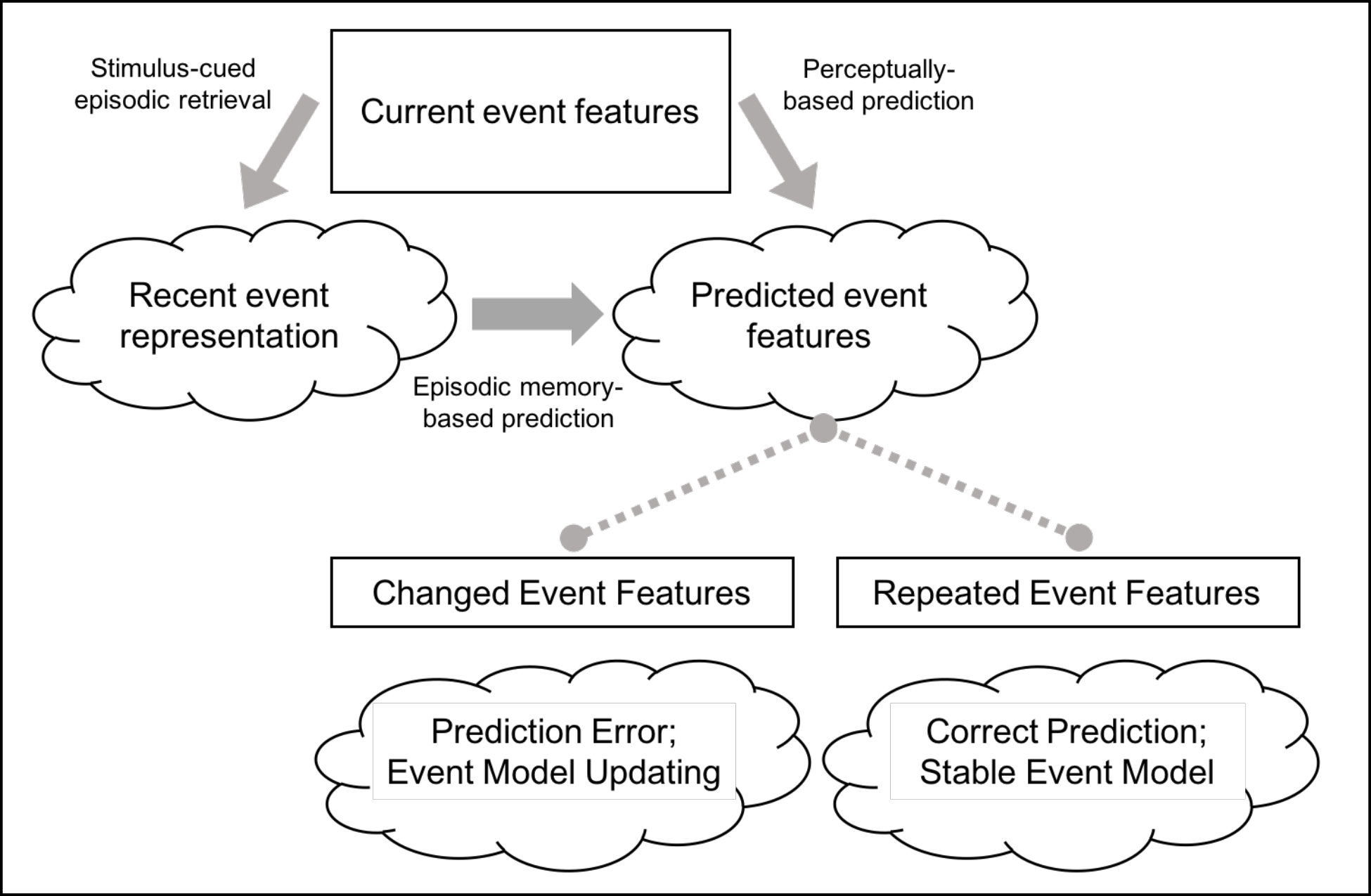
Schematic depiction of the processing chain leading to episodic change detection according to Change Comprehension Theory. Boxes represent perceived features and clouds represent cognitive representations. Arrows indicate the flow of information from perceptual inputs and memory representations to predictions. Dashed lines indicate the influence of predictions on the perception of upcoming event features. The theory proposes that current event features cue retrieval of recent related event representations. Both those representations and ongoing perceptual information inform predictions about upcoming event features. Changed features in upcoming events lead to prediction error and event model updating, whereas repeated features tend to lead to maintaining stable event models.

Figure 1 provides a schematic depiction of the proposed processes involved in detection of event changes according to CCT. The account proposes that as people observe everyday events, they are constantly predicting upcoming events that will occur in the near future. During these observations, people encounter activity features that cue recollection of episodic representations of recent related events. These recollections can occur involuntarily as a consequence of feature overlap between current and earlier events (Berntsen, 1996; Hintzman, 2011) or through self-initiated elaboration of event features as retrieval cues (Jacoby, 1974; Jacoby & Wahlheim, 2013). Regardless of how they are accomplished, recollections of earlier events should affect observers' predictions about what will happen in the new situation. Most of the time, retrieval of episodic memories should improve prediction, because natural activity often contains repeating sequences of features. In these cases, accurate predictions would maintain stable event models. However, when features change from one event to another related event, this can lead to a prediction error that upregulates attention to unexpected features leading to change detection and event model updating.

In the short term, episode-based prediction errors are likely to incur a processing cost and to interfere with ongoing processing. However, these very costs may have long-term benefits for episodic memory encoding, if they allow the observer to encode a representation that includes: (a) the original prediction of a repeated event, (b) the fact of the prediction error, and (c) the unexpected features. The formation of configural representations that comprise changed features and the cognitive operations associated with change detection will allow the observer to subsequently remember what happened during the original event and the new event, and the temporal relationship between the two. This will produce proactive facilitation of the original event for retrieval of the new event when the configural trace can be accessed via change recollection. However, when changes cannot be recollected, proactive interference should occur, as the retrieval of the original event that allowed for change detection should increase the accessibility of its competitive features.

An important aspect of CCT is that the memorial benefits of detecting and recollecting change depend on the cognitive system registering (consciously or not) the prediction error. If two events are simply encoded as separate instances when change is not detected, one would expect proactive interference rather than facilitation to the extent that episodes belong to similar event models and compete during retrieval (Radvansky, 2005, 2012). However, when change is not detected, two related events belonging to distinct event models would be less competitive, thus reducing interference. Such effects of event segregation are reminiscent of research showing reductions in interference due to context differentiation (for reviews, see Abra, 1972; Smith & Vela, 2001). However, although perfect context differentiation should eliminate proactive interference, it cannot produce proactive *facilitation*, because memory for independent traces cannot exceed that of memory for control events that only occurred once.

In addition to providing a mechanistic account of the processes involved in the comprehension of event changes, CCT also has implications for understanding age-related differences in the comprehension of such changes. Older adults experience deficits in attention and episodic memory (for reviews, see Balota et al., 2000; Zacks et al., 2000) that affect their ability to bind event features (e.g., Naveh-Benjamin, 2000) and perceive structure in ongoing activities (Kurby & Zacks, 2011; Zacks et al., 2006). According to CCT, these deficits should lead to poorer encoding of discrete event representations. The diminished coherence of event representations would reduce the frequency of stimulus-cued retrievals of prior events due to fewer perceived overlapping features with current events. Consequently, older adults should experience fewer episode-based prediction errors than younger adults leading to the detection and recollection of fewer event changes. When older adults do detect changes, their diminished ability later recollect changes (Wahlheim, 2014) should lead them to access fewer configural representations, resulting in a greater susceptibility to interference among competing activities features. If this proposal is correct, specific impairments in aspects of this processing chain may be targeted for remediation.

## The Present Experiments

To provide initial tests of our proposal, we developed the *everyday changes* paradigm, which combines procedures from the event perception and episodic memory literatures. In this paradigm, older and younger adults viewed movies depicting an actor performing everyday activities across the course of two fictive days in her life. The relationship of stimulus features between these episodes are varied such that some activities recur with a critical feature being repeated (e.g., ordering a *cheeseburger* on both days), other activities have features that change across the days (cheeseburger/salad), and control activities only appear on the second day (salad only). This design parallels the verbal learning A-B, A-D design, including that used by Jacoby, Wahlheim, and colleagues (Jacoby et al., 2015; Wahlheim & Jacoby, 2013). However, as mentioned above, natural activities differ from verbal paired associates in an important way: Natural activities have meaningful temporal structure. For example, the association between entering a restaurant and ordering a meal is a reliable regularity that is learned over many experiences with restaurant meals. The everyday changes paradigm captures this important feature of natural activities. After viewing both movies, participants are given a final cued-recall test on which they are asked to: 1) recall Day 2 activity features, 2) indicate whether features of the activity changed between days, and 3) recall Day 1 activity features for the activities that they remember changing between days.

We derived our hypotheses from the findings in earlier studies showing that detection and recollection of change counteracts interference, presumably through the encoding and retrieval of configural traces comprised of multiple event features and the cognitive operations that associate those features (e.g., Jacoby et al., 2015; Wahlheim & Jacoby, 2013). We hypothesized that cued recall of Day 2 features would show proactive facilitation when participants detected and recollected changed features, and proactive interference when participants detected but did not recollect changed features. Conversely, we expected that detecting but not recollecting changed features would lead to more intrusions of Day 1 features due to the retrieval-induced boost in accessibility from retrieval practice, whereas detecting and recollecting changes would reduce those errors due to the benefits on order memory conferred by accessing configural traces. Critically, we propose that the change detection necessary for encoding configural traces results from episode-based prediction errors.

We further hypothesized that older adults would be more susceptible to interference, and this would reflect poorer detection and recollection of change relative to younger adults. As described above, we derived this hypothesis from the fact that older adults have well-established deficits in attention and memory (for reviews see, Balota et al., 2000; Zacks et al., 2000) that extend to the abilities to form associations among features of perceived stimuli (e.g., Naveh-Benjamin, 2000) and to perceive and remember structure in ongoing events (Kurby & Zacks, 2011; Zacks et al., 2006). These deficits should lead to impaired encoding of event features, which would reduce older adults' ability to detect change. This would result from impoverished event representations decreasing the feature overlap necessary for current events to cue recall of recent related events. Older adults should then be *more* accurate in their predictions for changed activity features because they would be less likely to use the previous event to predict features of the new event. However, even when older adults do make contact between activities that lead to prediction errors, their binding deficit could diminish the formation of configural traces, rendering them less likely to represent together the retrieval event, the fact of the prediction error, and the temporal relationship between events. This could result in older adults being less likely to recall both related events when recollecting change. We describe specific hypotheses for each of the two present experiments below.

## Experiment 1

Experiment 1 examined the effects of episodic changes on memory for features of naturalistic activities. Participants first viewed two movies of an actor performing everyday activities on two fictive days in her life. The activities were either repeated, changed, or new (control) on Day 2. One week later, participants returned for a second session and took a cued recall test that asked about the activities of Day 2 and also whether each activity had changed from Day 1 to Day 2—a judgment of *change recollection*. Based on our proposal described above, we hypothesized successful change recollection would indicate that the participant had experienced a prediction error during the viewing of the changed activity during the Day 2 movie, resulting in updating and the formation of a configural representation of Day 2 that included the retrieval of the Day 1 and the ensuing prediction error as a component. Therefore, we hypothesized that successful change recollection would be associated with better recall of Day 2 activity features and lower rates of intruding Day 1 features. Based on earlier findings showing that older adults are more susceptible to interference effects (Healey, Hasher, & Campbell, 2013; Jacoby, Debner, & Hay, 2001; Wahlheim, 2014), we expected older adults to show a differential deficit for recall of changed activities, which would partly reflect a change recollection deficit.

## Method

The full stimulus sets for all the materials that we used in the present experiments (i.e., movies of the actor performing everyday activities and cued recall questions), anonymized data files, data with coded responses, and analysis scripts can be downloaded from the following URL: https://osf.io/t2qjk/. The research reported here was by the Institutional Review Boards of Washington University in St. Louis (Experiments 1 and 2) and the University of North Carolina at Greensboro (Experiment 2).

### Participants

The participants were 36 younger adults (26 females; *M*_Age_ = 20.19, *SD* = 1.43, range: 18-23) from Washington University in St. Louis and 36 older adults (27 females; *M*_Age_ = 77.19, *SD* = 8.07, range: 61-94) from the St. Louis community. Three older adults were excluded from analyses because they scored lower than 25 on the Mini-Mental State Exam (MMSE; Folstein, Folstein, & McHugh, 1975). The final sample of 33 older adults (25 females; *M*_Age_ = 77.00, *SD* = 7.79; range: 61-90) all had MMSE scores of 25 or above (*M* = 28.61, *SD* = 1.60, range: 25-30). The overall sample size was determined from power analyses conducted in G×Power Version 3.1.9.2. We based our sample size on an earlier study examining proactive effects of memory and the moderating effects of change recollection in younger and older adults (Wahlheim, 2014).

The effect sizes from those experiments showing significant main effects of age on cued recall performance were large. We converted the smallest of those effect sizes (η_p_^2^ = .20) to Cohen’s *f* = 0.5. A minimum sample size of 17 per group was necessary to detect comparable effects with power = .80 and α = .05. We chose a larger sample size here because we knew less about how the current materials would behave. Younger adults were either paid $10 per hour or were given partial course credit, and older adults were paid $10 per hour. Vocabulary scores on the Shipley Institute of Living Scale (Shipley, 1986) were significantly higher for the 33 older adults in the final sample (*M* = 36.33, *SD* = 2.70) than for the 30 younger adults who took the vocabulary test (*M* = 34.67, *SD* = 2.26), *t*(61) = 2.64, *p* = .01, *d* = 0.67. (Six younger adults did not take the vocabulary test due to experimenter error.)

### Design and Materials

A 2 × 3 mixed factorial design was used. Age (younger vs. older) was a between-subjects variable, and Activity Type (repeated, control, or changed) was manipulated within-subjects. The materials were videos of a female actor performing daily activities on two fictional days in her life. The activities took place in or around the actor’s home. There were two versions of each activity that differed on a thematically central feature (see Figure 2). For some of the activities, the changed feature was an object that the actor interacted with (e.g., the towel she hung in the bathroom; Figure 2, top panels), whereas for other activities, the changed feature was the action itself (e.g., the type of exercise she performed on a yoga mat; Figure 2, bottom panels).

**Figure 2.**
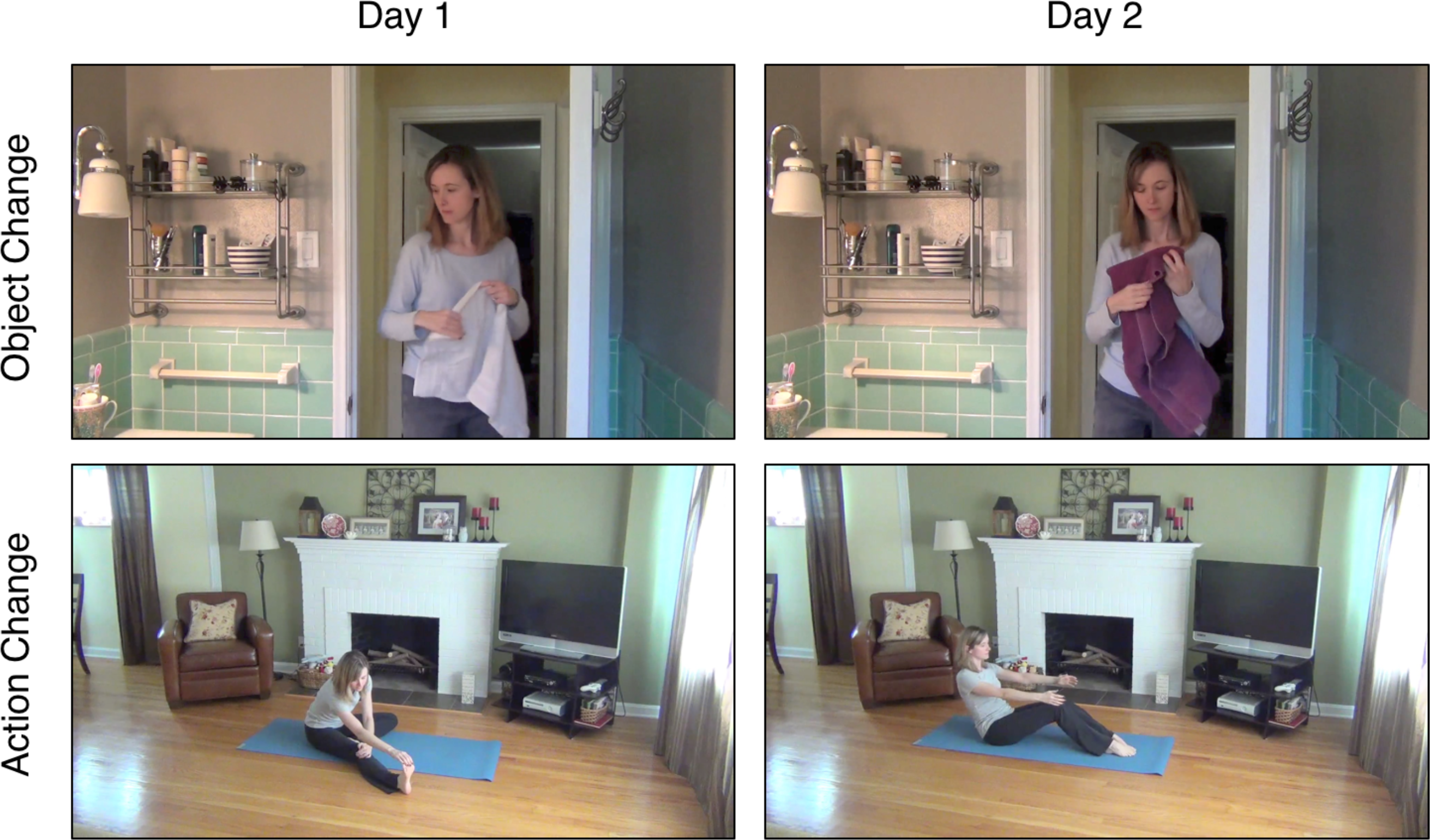
Example still frames from changed activities. The left column shows examples of activities from a Day 1 movie, and the right column shows examples of changed versions of those activities that appeared in the Day 2 movie. The top row shows an example of an object change and the bottom row shows an example of an action change.

The material set consisted of 62 total activities (51 critical; 11 untested fillers). For the critical activities, 45 included object changes and 6 included action changes. For the filler activities, 10 included object changes and 1 included an action change. To counterbalance activities across activity types, the 51 critical activities were divided into 3 groups of 17. Each activity appeared once in each activity type condition across three experimental formats. To create the activity type conditions, we varied the versions of activities in the Day 1 movie across three formats, leaving out control activities that only appeared in Day 2, and held constant the version of activities in the Day 2 movie.

Each Day 1 movie format depicted the actor performing 45 activities (34 critical; 11 untested fillers). The durations of the Day 1 movie formats were: 29 min and 15 s (Format 1), 30 min and 14 s (Format 2), and 28 min and 37 s (Format 3). The Day 2 movie depicted the actor performing 62 activities (51 critical; 11 untested fillers); its duration was 41 min and 24 s. The activity type conditions appeared in fixed-random orders such that no more than three subsequent critical activities were from the same condition in any movie. The average serial position of activities in each movie was equated across critical activity types to control for serial position effects on memory. Untested filler activities were interspersed among the critical activities to improve the coherence of action sequences. All fillers were repeated across movies.

Sixty-two activity test cues appeared on the final test. Each cue probed participants’ memory for the central feature of the relevant activity (e.g., “What type of towel did the actor hang in the bathroom?”).

### Procedure

Participants completed tasks in two sessions separated by approximately one week. The average delay between sessions (*M* = 7.10 days, *SD* = 0.63, range: 6 – 12) did not differ between older and younger adults. All movies were presented at a 1280 × 720 aspect ratio. The instructions, movies, and memory test described below were all presented using E-Prime 2 software (Schneider, Eschman, & Zuccolotto, 2002).

During Session 1, participants watched an actor perform everyday activities on two fictive days in her life. Prior to each movie, participants were told to pay attention to the activities. Before watching the second movie, participants were also told that the upcoming activities took place later in the actor’s week. Participants were not informed about the how activities were related between days.

During Session 2 (one week later), participants were given a cued recall test for Day 2 activity features that included change recollection judgments. Test cues appeared in the same order as the original Day 2 activities to create approximately equal retention intervals for each. Before the test began, participants were shown two clips of example activities that depicted the type of change that participants were supposed to report remembering. In both example activities, the actor wiped down her kitchen counter. The changed feature was the implement that she used to clean the counter (*wash cloth* or *paper towel*). When participants indicated that an activity had changed on the test, they were asked to recall the Day 1 feature. For each cue, a question appeared asking participants about the criterial Day 2 feature (e.g., “What type of towel did the actor hang in the bathroom?”), and participants typed their responses onto the screen (e.g., “maroon bath towel”). After responding, participants were asked whether the way the actor accomplished the activity changed from Day 1 to Day 2. Participants pressed the “1” key to indicate that the activity had changed and the “2” key to indicate that the activity had not changed. When participants indicated that the activity had changed, they were asked to type the Day 1 feature (e.g., “white hand towel”) onto the screen. Participants proceeded to the next question after making this response or after indicating that no change had occurred. Most participants typed their own responses, but some older adults preferred to have the experimenter type for them.

After the recall test, all participants completed a computerized version of the Shipley vocabulary test. The MMSE was administered to older adults at the end of the session.

## Results and Discussion

The level for significance was set at *α* = .05. Participants’ responses when attempting to recall Day 2 features were classified into four types; we will illustrate these by considering a case in which the actor hung a *white hand towel* on Day 1 and a *maroon bath towel* on Day 2 during the corresponding activity. *Day 2 recall* responses were correct descriptions of the criterial activity feature in the Day 2 movie (“she hung a maroon towel”); *Day 1 intrusions* were responses that included the criterial feature from Day 1 (“she hung a small white towel”) instead of the Day 2 feature; *Ambiguous* responses were correct action descriptions that did not distinguish between days (“she hung a towel”); and *Other Error* responses were either commission errors that did not include criterial features from the target activity on either day or were omitted responses. Note that commission errors in this class of responses sometimes included criterial features from non-target activities and other times included features that were not associated with any earlier-viewed activity. We did not have any a *priori* hypotheses regarding Ambiguous and Other Error responses, so we do not report analyses of those responses here. Two raters independently coded responses. Discrepancies were resolved through discussion. Cohen’s *κ* for the initial ratings (*κ*; = .87, *p* < .001) showed almost perfect agreement between raters (Landis & Koch, 1977).

To examine how age and the viewing of Day 1 activities affected memory for Day 2 activities, we computed response probabilities for Day 2 recall and Day 1 intrusions for each activity type separately for younger and older adults. We compared these probabilities for repeated and changed activities with control activities using separate logistic mixed effects models, with Age and Activity Type as fixed effects, and subjects and activities as random effects. Modeling the random effect of activities was especially important when examining conditional probabilities in recall due to the potential for item selection effects. Mixed effects models were fitted using the lme4 package in R (Bates, Maechler, Bolker, & Walker, 2015), hypothesis tests were performed using the Anova function of the car package (Fox & Weisberg, 2011), and post hoc comparisons using the Tukey method were conducted using the lsmeans package (Lenth, 2016). The logistic models operate on log-likelihoods; for data presentation we have converted estimates of cell means and confidence intervals back to probabilities.

## Recall Performance

**Figure 3.**
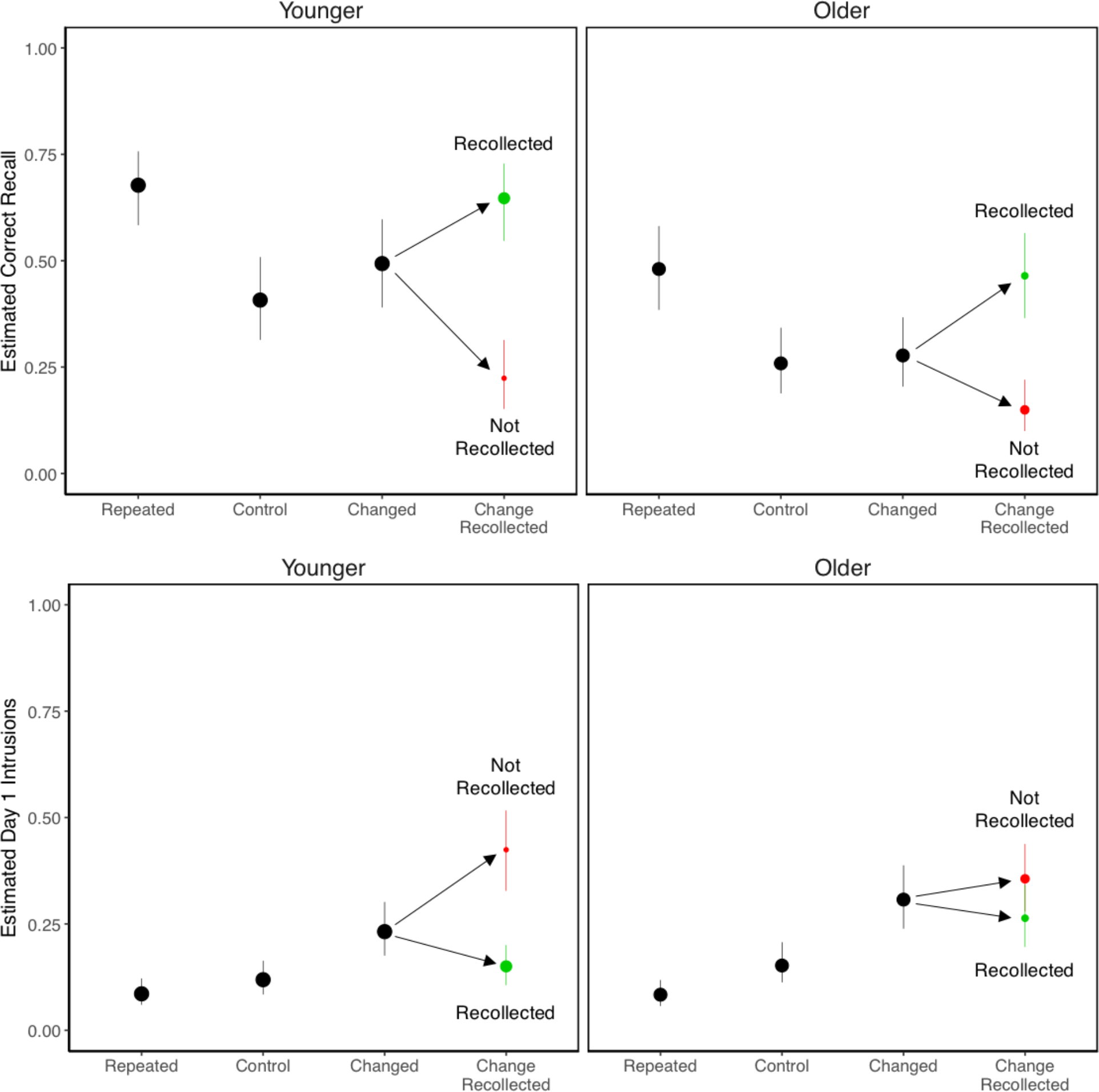
Probabilities of Day 2 correct recall (top panels) and Day 1 intrusions (bottom panels). Conditional points (in green and red) refer to change recollection probabilities. Green points indicate that change was recollected and red points indicate that change was not recollected. Conditional point areas represent proportions of observations contributing to each cell. Error bars are bootstrap 95% confidence intervals.

Day 2 recall performance was computed for critical activities only. The analysis of Day 2 recall (Figure 3, top panels) revealed that younger adults recalled significantly more activities than older adults, χ^2^(1) = 24.01, *p* < .001. In addition, repeated activities were remembered best and control activities were remembered worst, leading to a significant main effect of Activity Type, χ^2^(2) = 124.31, *p* < .001. Post hoc tests showed that, for both age groups, all three activity types differed from each other, smallest *z* ratio = 2.64, *p* = .02, with one exception: For older adults, the changed and control activities were not significantly different, *z* ratio = 0.66, *p* = .79. Despite this, the Age × Activity Type interaction was not significant, χ^2^(2) = 1.69, *p* = .43.

The analyses of Day 1 intrusions (Figure 3, bottom panels) revealed that older adults produced more Day 1 intrusions than younger adults, leading to a significant main effect of Age, χ^2^(1) = 4.34, *p* = .037. In addition, both age groups produced Day 1 intrusions most often for changed activities and least often for repeated activities, leading to a significant main effect of Activity Type, χ^2^(2) = 144.38, *p* < .001. For repeated and control activities, Day 1 intrusions tended to be either participants misremembering the actor interacting with objects that were visible in the scene but were not the criterial feature, or participants guessing the alternative object or action despite not having seen that version on either day. The Age × Activity Type interaction was not significant, χ^2^(2) = 3.09, *p* = .21. Post hoc tests showed that, for younger adults, all three activity types differed significantly from each other, smallest *z* ratio = 3.76, *p* < .001, and for older adults, changed activities differed significantly from control and repeated activities, smallest *z* ratio = 5.23, *p* < .001, and control activities differed marginally from repeated activities, *z* ratio = 2.06, *p* = .098.

## Change Recollection

Table 1 displays the probabilities of participants indicating on the cued recall test that activities had earlier changed from Day 1 and Day 2. The probability of correct change recollection for changed activities was greater than the probabilities of false change recollection for repeated and control activities, leading to a significant main effect of Activity Type, χ^2^(2) = 462.94, *p* < .001. Younger adults indicated that more activities changed than older adults, leading to a significant main effect of Age, χ^2^(1) = 10.91, *p* < .001. A significant Age × Activity Type interaction indicated that this difference was greatest for changed activities, χ^2^(2) = 21.60, *p* < .001. These results showed more accurate change recollection for younger than older adults.

**Table 1.**
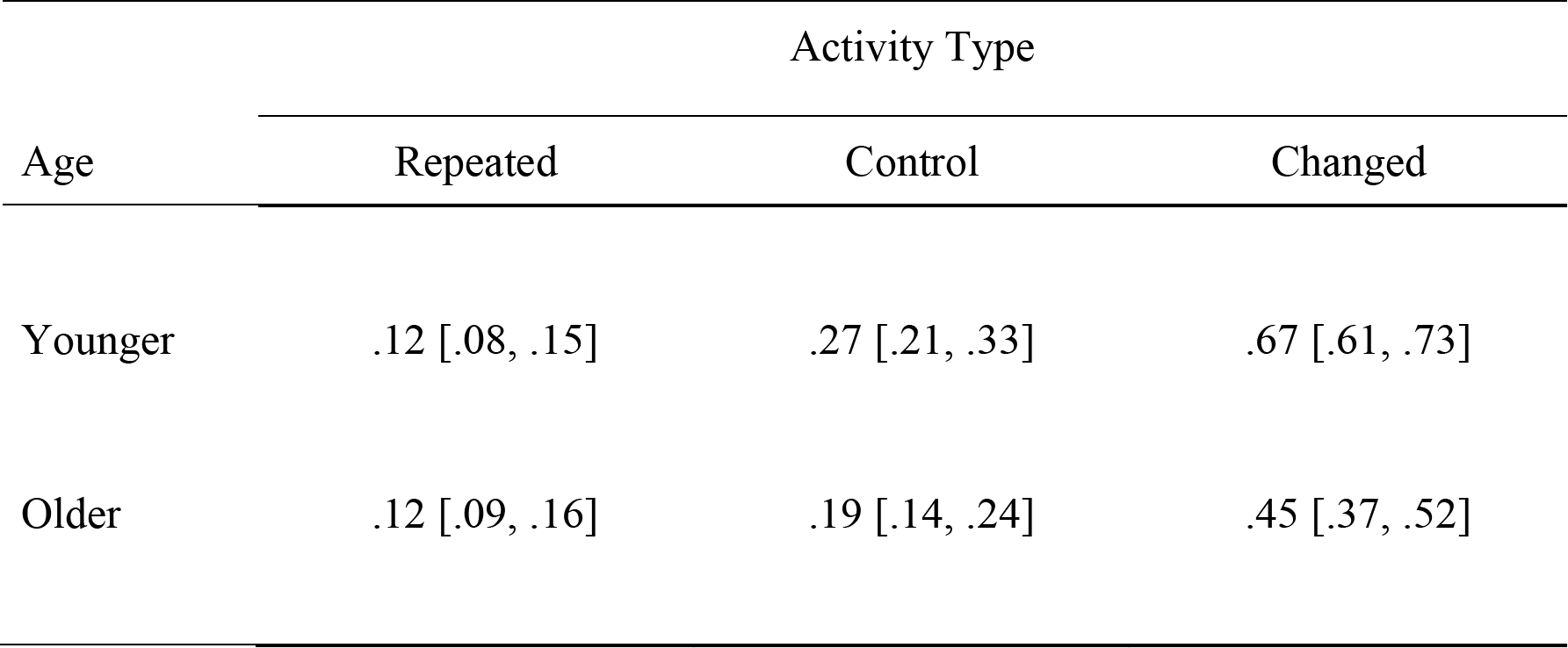
Probabilities of Change Recollection as a Function of Age and Activity Type: Experiment 1

Note: Probabilities for Changed activities are correct classifications, whereas probabilities for Repeated and Control activities are incorrect classifications. Bootstrap 95% confidence intervals are displayed in brackets.

## Correct Day 1 Recall Following Change Recollection

When participants indicated that activities had changed between Day 1 and Day 2, they also were asked to recall the original Day 1 activity features. We included this measure to assess the quality of configural representations that were established during Day 2 viewing. We coded these responses in the same manner as for Day 2 recall. Two independent raters showed almost perfect agreement in their ratings (*κ* = .85, *p* < .001). Consistent with our predictions, Day 1 recall following change recollection was significantly lower for older (*M* = .55, *CI* = [.44, .66]) than for younger (*M* = .73, *CI* = [.65, .81]) adults, χ^2^(1) = 7.68, *p* < .001. These results showed that configural traces were of lower quality for older than younger adults.

## Recall of Changed Activities Conditionalized on Change Recollection

As described in the Introduction, we expected that correct change recollection would be associated with enhanced memory for Day 2 activity features. This prediction is based on the assumption of CCT that change recollection should allow participants to retrieve activity features from both days along with the cognitive operations that occurred during encoding, which should facilitate memory for temporal order. To test this assumption, we examined recall of changed activities conditionalized on change recollection (red and green points in Figure 3), we fit separate logistic mixed effects models to Day 2 recalls and Day 1 intrusions, with Age and Change Recollection as fixed effects, and subjects and activities as random effects. We do not report redundant main effects of Age here.

The memorial benefits of change recollection were examined separately for Day 2 recall and Day 1 intrusions by conditionalizing performance on whether change was recollected. Based on our earlier predictions, these benefits should take the form of increased Day 2 recall and decreased Day 1 intrusions. The analysis of Day 2 recall revealed better memory when change was recollected than when it was not, leading to a significant main effect of Change Recollection, χ^2^(1) = 111.11, *p* < .001. There was no significant Age × Change Recollection interaction, χ^2^(1) = 0.65, *p* = .42. Post hoc tests comparing Day 2 recall of changed activities conditionalized on change recollection to control activities were conducted next to examine proactive effects of memory. For both age groups, Day 2 recall was significantly higher for changed than control activities when change was recollected, *smallest z* ratio = 4.33, *p* < .001, whereas recall was significantly lower for changed than control activities when change was not recollected, *smallest z* ratio = 6.34, *p* < .001. These results were consistent with earlier findings showing proactive facilitation resulting from change recollection and proactive interference in the absence of change recollection.

The analysis of Day 1 intrusions revealed fewer intrusions when change was recollected than when it was not, leading to a significant main effect of Change Recollection, χ^2^(1) = 36.86, *p* < .001. Change recollection reduced Day 1 intrusions more for younger than older adults, resulting in a significant Age × Change Recollection interaction, χ^2^(1) = 11.73, *p* < .001, which could reflect age-related deficits in the binding of features within configural traces.

We also examined whether the memorial benefits of change recollection depended on participants being able to recall Day 1 activity features. The notion here is that being able to recollect change along with both activity features may be necessary to obtain the benefits of retrieving configural traces. To examine this, we calculated Day 2 recall conditionalized on three levels of change recollection: change recollected and correct recall of Day 1 features, change recollected without recall of Day 1 features, and change not recollected (Table 2, top rows). We fit these data with a logistic mixed effects model that included Age and Change Recollection (recollected with Day 1 recall, recollected without Day 1 recall, and not recollected) as fixed effects and subjects and activities as random effects.

**Table 2.** Day 2 Recall conditionalized on Change Recollection and Day 1 Recall as a Function of Age: Experiments 1 and 2

| Experiment | Age | Change Recollected |  | Change Not Recollected |
| --- | --- | --- | --- | --- |
|  |  | Day 1 Recalled | Day 1 Not Recalled |  |
| Experiment 1 | Younger | .84 [.77, .91] | .20 [.10, .29] | .20 [.12, .28] |
|  | Older | .71 [.59, .83] | .22 [.12, .32] | .14 [.08, .20] |
| Experiment 2 | Younger | .86 [.80, .93] | .33 [.19, .47] | .40 [.29, .51] |
|  | Older | .64 [.51, .77] | .30 [.17, .43] | .26 [.17, .34] |

Note: Bootstrap 95% confidence intervals are displayed in brackets.

Note: Bootstrap 95% confidence intervals are displayed in brackets.

When change was recollected, Day 2 recall was higher when Day 1 features were recalled relative to when they were not, leading to a significant main effect of Change Recollection, χ^2^(2) = 196.33, *p* < .001. There was no significant Age × Change Recollection interaction, χ^2^(2) = 4.25, *p* = .12. Post hoc tests revealed no significant difference in Day 2 recall when change was recollected without recall of Day 1 features, and when change was not recollected, largest *z* ratio = 1.78, *p* = .18. These results show that enhanced Day 2 recall resulting from change recollection depended on the ability to correctly recall Day 1 activity features. This finding suggests that the facilitative effects of change recollection might only be obtained when the original activity features are accessible at retrieval.

## Summary

The results from Day 2 recall performance showed that repeated viewing of the same event increased performance relative to viewing an event once, and that changing a central feature of an event resulted in proactive facilitation when the change was recollected and proactive interference when the change was not recollected. In contrast, fewer Day 1 intrusions were produced when change was recollected than when it was not. The proactive facilitation in Day 2 recall was driven by instances when change was recollected and Day 1 features were recalled. CCT predicts this, because Day 2 recall should be facilitated by forming a configural trace that also includes the retrieval of Day 1 features during viewing of Day 2. According to CCT, a crucial stage leading to having a configural trace available during final recall is experiencing a prediction error due to the changed feature during Day 2 viewing. However, this experiment did not include a direct measure of change *detection*, only change recollection, so we were unable to examine the role of change detection in establishing such configural representations. Experiment 2 was conducted to obtain such measurements, while also attempting a replication of the findings of Experiment 1.

### Experiment 2

Experiment 1 showed proactive facilitation in memory for changed activity features when change was recollected for both age groups. This finding is consistent with earlier studies (e.g., Jacoby et al., 2015; Wahlheim, 2014) as well as our *a priori* predictions. However, Experiment 1 did not directly examine the combined effects of detecting and recollecting episodic changes in ongoing activities. To address this, we modified the everyday changes paradigm in Experiment 2 by including a change detection measure that followed the presentation of each individual activity. Pilot testing showed that change detection performance was on ceiling when movies from both fictive days were shown consecutively in the same session, so we inserted a one-week delay between presentations of Day 1 and Day 2 movies. We also included a measure of change recollection on the final recall test that was given one week following Day 2 movies. In contrast to Experiment 1, the measures of change detection and change recollection both included three possible classification responses that corresponded to each activity type. We made this change to assess participants’ awareness of the relationship between all Day 2 activities and the activities shown in the Day 1 movie. We also examined participants’ meta-awareness of the accuracy of their classifications by collecting confidence judgments. Older adults are typically impaired in their metacognitive monitoring of recollective information (for a review, see Dodson, 2017), and this allowed us to determine if this deficit extends to activity type classifications, which may often be recollective experiences.

Consistent with the predictions of CCT, we expected that detecting changes during Day 2 viewing would serve to encode features from Day 1 activities within representations that also included Day 2 features. Consequently, Day 2 recall should be enhanced when participants are able to access those traces using recollective processes. However, the retrieval practice that occurs during change detection should also increase the accessibility of Day 1 activity features, which would make them more competitive during retrieval on the final test when recollective processes cannot be deployed to oppose the accessibility of those features. This leads to the prediction that Day 2 recall will be enhanced when change is detected and recollected, and impaired when it is detected but not recollected. As another consequence of the opposing effects of recollection, Day 1 intrusions should be lower when change is detected and recollected, whereas intrusions should be higher when change is detected but left unopposed by change recollection. This trade-off should determine overall levels of recall performance, and also determine the degree of older adults’ differential deficit for changed activities.

Based on the fact that older adults have impaired encoding of event structure (Kurby & Zacks, 2011; Zacks et al., 2006), we hypothesized that older adults would make contact between related activities on each day less often than younger adults. This should result in older adults being less able to accurately detect and recollect changed activities as such and less able to remember Day 1 activity features when correctly classifying changed activities. This age-related deficit should then have detrimental consequences for Day 2 recall, which would in part explain older adults’ differential deficit. Finally, this constellation of age-related deficits might also have consequences for metacognitive awareness of classification accuracy. The classification of the relationships between activities on separate episodes should rely on recollective processes. In addition, older adults have often been shown to have metacognitive deficits when relying on recollective bases for retrieval and subsequent evaluations of memory accuracy (for a review, see Dodson, 2017). Thus, to the extent that activity type classifications are based on recollection, older adults should show poorer metacognitive accuracy in their confidence judgments.

## Method

### Participants

We initially planned to collect data from at least 36 participants per age group to match the sample sizes used in Experiment 1. We scheduled appointments for more than 36 younger adults anticipating that some may drop out before completing all three sessions. The entire sample of younger adults included 44 participants from Washington University in St. Louis. However, data from six younger adults were excluded from analyses because five participants dropped out before completing all three sessions, and the program crashed for one participant. The final younger adult data set included 38 participants (23 females; *M*_Age_ = 19.84, *SD* = 1.26, range: 18-23). For older adults, we recruited a total of 37 participants from the Greensboro, NC community. Data from one older adult were excluded because the program crashed before the experiment was completed. The final older adult data set included 36 participants (20 females; *M*_Age_ = 70.22, *SD* = 2.88, range: 65-75). All the older adults had MMSE scores of 25 or above (*M* = 28.44, *SD* = 1.25, range: 25-30). Younger adults were given partial course credit, and older adults were paid $10 per hour. Vocabulary scores were significantly higher for older (*M* = 34.97, *SD* = 3.44) than younger (*M* = 33.50, *SD* = 2.24) adults, *t*(72) = 2.19, *p* = .03, *d* = 0.51.

### Design, Materials, and Procedure

The design and materials were identical to Experiment 1, except that the Day 2 movies were presented as individual clips so that activity classifications could be made following each. The procedure maintained most of the key elements of Experiment 1, but we modified aspects related to the measurement of participants’ processing of change. Whereas in Experiment 1 we measured only participants’ *recollection* of change on the final recall test, in Experiment 2 we added a measure of their *detection* of change while viewing the Day 2 movie. This necessitated adding a one-week delay between the viewing of Day 1 and Day 2 movies so that there would not be a ceiling effect in change detection performance. Experiment 2 was administered in three sessions: 1) Day 1 movies during the first session, 2) Day 2 movies during the second session, and 3) cued recall during the third session. Appointments were scheduled with the intention of making the between-session intervals as close to one week as possible. The mean delay across sessions (*M* = 7.16 days, *SD* = 1.17, range: 6-17) did not differ between older and younger adults.

During Session 1, participants were instructed to pay attention as they watched the Day 1 activities. They were also shown the earlier-described example activity depicting the actor cleaning the kitchen counter with a wash cloth before viewing the Day 1 activities. This example was included so that it could be referenced in the change detection instructions on Session 2.

During Session 2, the Day 2 activities appeared as individual clips. The duration of each activity varied widely (*M* = 38.23 s, *SD* = 25.86 s, range: 4-109 s). Participants were instructed to pay attention to the activities and classify them based on their relationship to the Day 1 activities, which included detecting changes. To clarify the types of changes that should be detected, participants were shown the example of the actor cleaning the kitchen counter with a washcloth again with reference to the example activity from Session 1. Following this, participants watched the changed version of that activity depicting the actor cleaning the kitchen counter with a paper towel. When viewing the actual Day 2 activities, participants indicated how each individual clip related to the activities on Day 1. After each clip concluded, a prompt appeared asking whether activities were the same as Day 1, changed from Day 1, or new on Day 2. Participants indicated “same” by pressing the “1” key, “changed” by pressing the “2” key, and “new” by pressing the “3” key. After each response, participants rated how confident they were in their classification on a scale from 1 (not) – 5 (very) by pressing the corresponding key. When participants indicated that an activity had changed, they were asked to describe the Day 1 feature following their confidence judgment by typing their response onto the screen.

During Session 3, participants completed the cued recall test of Day 2 features, that also tested for change recollection. This test was similar to the test from Experiment 1, except that the activity type classifications included all three possible activity types. As with the classifications made in Session 2 of the present experiment, participants indicated “same” by pressing the “1” key, “changed” by pressing the “2” key, and “new” by pressing the “3” key. Most participants typed their responses, but some older adults preferred to have the experimenter type for them.

## Results and Discussion

### Recall Performance

We coded recall responses in the same manner as Experiment 1. Two raters initially showed substantial agreement in their classifications, *κ* = .67, *p* < .001; disagreements were resolved by discussion. We compared Day 2 recall and Day 1 intrusions for each activity type for younger and older adults (Figure 4) using the same logistic mixed effects models as in Experiment 1.

**Figure 4.**
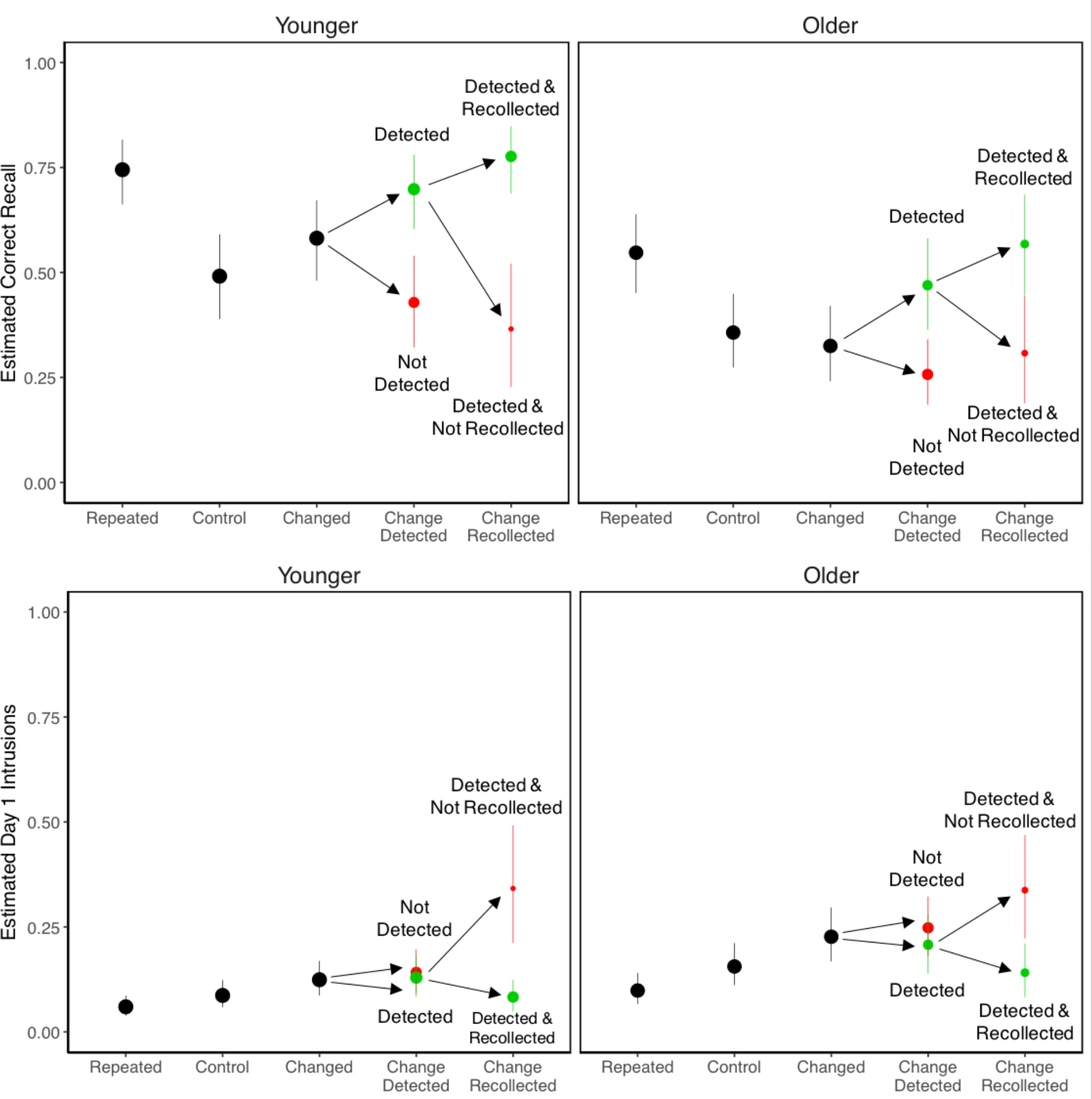
Probabilities of Day 2 correct recall (top panels) and Day 1 intrusions (bottom panels). Conditional points (in green and red) for changed activities refer to change detection and change recollection probabilities. Green points correspond to detected or recollected changes, whereas red points correspond to changes that were not detected or recollected. Conditional point areas represent proportions of observations contributing to each cell. Error bars are bootstrap 95% confidence intervals.

The analyses of Day 2 recall revealed higher performance for younger than older adults, leading to a significant main effect of Age, χ^2^(1) = 42.05, *p* < .001. Recall was best for repeated activities for both age groups, whereas recall of control activities was the lowest for younger but not older adults. This led to a significant main effect of Activity Type, χ^2^(2) = 115.45, *p* < .001, that was qualified by a significant Age × Activity Type interaction, χ^2^(2) = 7.70, *p* = .02. Post hoc tests indicated that recall differed across all three activity types, smallest *z* ratio = 2.86, *p* = .01, with one exception: For older adults, recall was not significantly different between changed and control activities, *z* ratio = 1.06, *p* = .54. These result show that older adults experienced a differential deficit in memory for changed activity features, as in Experiment 1. However, the statistical evidence is stronger in this case due to the significant interaction.

Older adults also produced more Day 1 intrusions than younger adults, leading to a significant main effect of Age, χ^2^(1) = 24.50, *p* < .001. Intrusions were produced most often for changed activities and least often for repeated activities, leading to a significant main effect of Activity Type, χ^2^(2) = 58.97, *p* < .001. The Age × Activity Type Group interaction was not significant, χ^2^(2) = 0.61, *p* = .74. Post hoc tests showed significantly more intrusions for changed than for repeated activities, smallest *z* ratio = 3.83, *p* < .001, whereas the difference between all other pairwise comparisons was marginally significant, smallest *z* ratio = 2.06, *p* = .099. The failure to replicate the significantly higher intrusion rate for changed than control activities in Experiment 1 was likely due to the longer interval between movies in Experiment 2.

### Detection and Recollection of Change

We excluded four older adult participants from the following analyses because inspection of their data suggested that they did not understand the activity type classification instructions. One participant appeared to have mis-mapped the response keys, confusing changed and control activities, whereas the other three participants never used the “new” classification. Table 3 displays the probabilities of correct classifications for activity types in session 2 (top rows) and session 3 (bottom rows). Note that these data are displayed differently from the comparable analyses in Experiment 1 because including all three response options in Experiment 2 allowed us to assess correct classification for each activity type. The data from each session were fit with separate logistic mixed effects models with Age and Activity Type as fixed effects and subjects and activities as random effects.

**Table 3.** Probabilities of Correct Activity Type Classification: Experiment 2

| Measure | Age | Activity Type |  |  |
| --- | --- | --- | --- | --- |
|  |  | Repeated | Control | Changed |
| Session 2 (Detection) | Younger | .82 [.79, .86] | .86 [.83, .89] | .58 [.52, .63] |
|  | Older | .74 [.69, .79] | .63 [.58, .69] | .43 [.38, .49] |
| Session 3 (Recollection) | Younger | .81 [.76, .85] | .59 [.53, .65] | .54 [.48, .60] |
|  | Older | .67 [.61, .73] | .35 [.29, .42] | .40 [.34, .47] |

Note: Bootstrap 95% confidence intervals are displayed in brackets.

Note: Bootstrap 95% confidence intervals are displayed in brackets.

The analyses of classification accuracy during session 2 showed that younger adults classified activity types more accurately than older adults, leading to a significant main effect of Age, χ^2^(1) = 51.62, *p* < .001. There was also a significant effect of Activity Type, χ^2^(2) = 225.86, *p* < .001, and a significant Activity Type × Age interaction, .^2^(2) = 18.35, *p* < .001. Post hoc tests showed that for younger adults, correct classification of repeated and control activities was not significantly different, *z* ratio = 1.82, *p* = .16, whereas correct classification was significantly higher for repeated and control than changed activities, smallest *z* ratio = 9.36, *p* < .001. For older adults, correct classification was significantly higher for repeated than control and for control than changed activities, smallest *z* ratio = 3.71, *p* < .001.

The analyses of classification accuracy during session 3 also showed that younger adults classified activity types more accurately than older adults, leading to a main effect of Age, χ^2^(1) = 43.80, *p* < .001. There was a significant main effect of Activity Type, χ^2^(2) = 206.19, *p* < .001, showing that classification accuracy was significantly higher for repeated than control and changed activities, smallest *z* ratio = 12.98, *p* < .001, and not significantly different between control and change activities, *z* ratio = 0.23, *p* = .97. The Activity Type × Age interaction was marginally significant, χ^2^(2) = 5.46, *p* = .06.

Together, the analyses from sessions 2 and 3 showed that older adults detected and recollected fewer changed activities than younger adults. These results also showed that older adults were generally impaired in their ability to classify activity types. We did not make any *a priori* predictions about whether these deficits would differ across activity types, so we do not have a clear interpretation for the interactions obtained above.

### Correct Day 1 Recall Following Change Recollection

When participants indicated that an activity changed between days on the final recall test (session 3), they were asked to subsequently recall the original Day 1 feature. Similar to Experiment 1, we used this measure to assess the quality of configural representations. Responses were coded in the same manner as Experiment 1. The raters showed almost perfect agreement in their initial ratings (*κ* = .81, *p* < .001). As predicted, older adults’ episodic memory deficit resulted in their recalling fewer Day 1 features after recollecting changes on the final test. Correct Day 1 recall following change recollection judgments was significantly lower for older (*M* = .51, [*CI* = .41, .62]) than younger (*M* = .75, [*CI* = .68, .83]) adults, χ^2^(1) = 18.38, *p* < .001. These results replicate Experiment 1 in showing poorer quality configural traces for older adults.

### Recall of Changed Activities Conditionalized on Change Detection and Recollection

As described above, we expected that detection and recollection of changed Day 2 activities would lead to enhanced memory their features. This was based on the assumption that change recollection should provide access to the contents of configural traces resulting in proactive facilitation of Day 2 recall. In the absence of change recollection, the retrieval involved in change detection should make Day 1 responses more accessible, resulting in those responses being more competitive with recall of Day 2 activity features. To test this, we conditionalized recall performance on detection and recollection of change (red and green points in Figure 4). We fit logistic mixed effects models to Day 2 recalls and Day 1 intrusions with Change Detection, Change Recollection, and Age as fixed effects, and subjects and activities as random effects. We do not report redundant main effects of Age here.

The analyses of Day 2 recall (Figure 4, top panels) revealed significant main effects of Change Detection, χ^2^(1) = 10.61, *p* = .001, and Change Recollection, χ^2^(1) = 25.02, *p* < .001, as well as a significant Change Detection × Change Recollection interaction, χ^2^(1) = 10.65, *p* = .001. The interaction showed that change recollection moderated the effects of change detection. Specifically, Day 2 recall was higher when change was detected and recollected relative to when change was detected but not recollected, *z* ratio = 6.26, *p* < .001. In contrast, when change was not detected, memory performance did not differ depending on whether change was recollected, *z* ratio = 1.01, *p* = .31. Consequently, we collapsed Day 2 recall when change was not detected across change recollection outcomes for presentation purposes in Figure 4. These effects were comparable for both age groups, leading to no significant interactions including Age as a variable, largest χ^2^(1) = 132, *p* = .25.

Post hoc tests comparing Day 2 recall of changed activities when change was detected conditionalized on change recollection to control activities were conducted next to examine proactive effects of memory. For both age groups, Day 2 recall was significantly higher for changed than control activities when change was recollected, smallest *z* ratio = 3.36, *p* = .002, whereas recall was significantly lower for changed than control activities when change was not recollected for older adults, *z* ratio = 2.59, *p* = .026, but not younger adults, *z* ratio = 1.76, *p* = .183. These results were consistent with earlier findings showing proactive facilitation resulting from change recollection for both age groups and proactive interference in the absence of change recollection for older adults. The finding that younger adults did not show proactive interference in the absence of change recollection may have resulted from too few observations to detect differences. However, the temporal separation between movies (one week) could have reduced the competition between event representations relative to Experiment 1. We return to this issue in the General Discussion.

The analyses of Day 1 intrusions (Figure 4, bottom panels) revealed no significant effect of Change Detection, χ^2^(1) = 197, *p* = .16, a significant effect of Change Recollection, χ^2^(1) = 23.68, *p* < .001, and a significant Change Detection × Change Recollection interaction, χ^2^(1) = 9.74, *p* = .002. As for Day 2 recall, this interaction showed that change recollection moderated the effects of change detection. When change was detected, there were significantly fewer Day 1 intrusions when change was recollected than when it was not recollected, *z* ratio = 4.68, *p* < .001. In contrast, when change was not detected, Day 1 intrusion rates did not differ depending on whether change was recollected, *z* ratio = 0.83, *p* = .41. Consequently, we collapsed Day 1 intrusions when change was not detected across change recollection outcomes for presentation purposes in Figure 4. These effects were comparable for both age groups, leading to a lack of significant interactions including Age as a variable, largest χ^2^(1) = 198, *p* = .15. These results show that change recollection counteracted Day 1 intrusions following change detection.

Similar to Experiment 1, we also examined whether the moderating effects of change recollection on change detection depended on correct Day 1 recall following change recollection. As described earlier, the idea was that recollecting change along with both activity features should confer the benefits of accessing configural traces. To examine this, we analyzed Day 2 recall of changed activities for which change was detected conditionalized on change recollection and Day 1 recall (Table 2, bottom rows). These data were fit with a mixed effects model that included Age and Change Recollection (change recollected and Day 1 recalled, change recollected and Day 1 not recalled, change not recollected) as fixed effects and subjects and activities as random effects.

This analysis revealed a significant effect of Change Recollection, χ^2^(2) = 113.35, *p* < .001, and a significant Age × Change Recollection interaction, χ^2^(2) = 6.30, *p* = .04. These results showed that the memory advantage in Day 2 recall when change was recollected was only obtained when Day 1 features were correctly recalled, and this advantage was greater for younger than older adults. This age difference could reflect higher configural trace quality for younger adults due to more effective change detection. For both age groups, correct Day 2 recall did not differ when change was recollected without Day 1 features and when change was not recollected, largest *z* ratio = 1.09, *p* = .52. Consistent with Experiment 1, these results show that the memorial benefits of change recollection depended on correct Day 1 recall. These results provide additional evidence for the assumption that detecting and recollecting change facilitates the encoding and retrieval of configural traces that protect against proactive interference.

### Confidence Judgments for Activity Type Classifications

**Figure 5.**
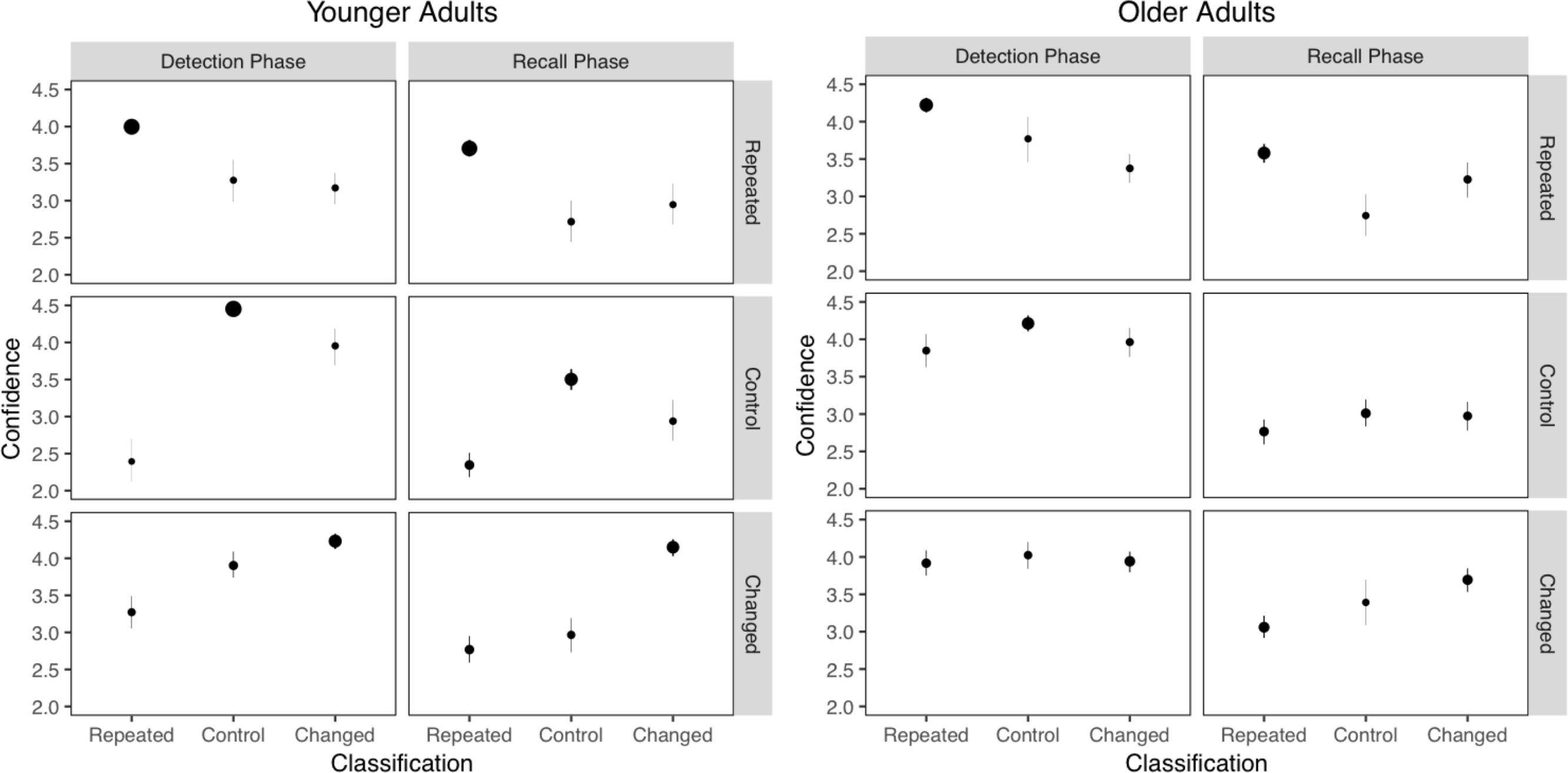
Mean confidence judgments as a function of age, activity type, and activity classification during session 2 (Detection Phase) and session 3 (Recall Phase) of Experiment 2. The point areas indicate the proportion of activities that were classified as each activity type. Error bars are bootstrap 95% confidence intervals.

Figure 5 displays the mean confidence judgments for activity type classifications on sessions 2 and 3. Our primary interest was whether participants’ confidence judgments discriminated between correct and incorrect classifications of activity types (i.e., monitoring resolution). Because individuals could use the confidence scales differently, we first z-scored each participant’s confidence judgments. Then, we submitted the confidence z-scores to multilevel linear models with Activity Type, Age, and Classification Accuracy as the fixed effects, and activity as a random intercept, and a random slope of accuracy by subject (to model the fact that monitoring resolution might vary by subject). In this analysis, better monitoring resolution is reflected in a larger effect of classification accuracy on confidence. That is, to the extent that one is aware of one’s classification accuracy, confidence should be higher for correct than incorrect classifications. Thus, one critical test is the main effect of accuracy on z-scored confidence. The other critical test is the interaction of accuracy with age, which indicates whether monitoring resolution differed between younger and older adults. We will report these two tests for classifications made in sessions 2 and 3.

Inspection of Figure 5 suggests that younger adults had good monitoring resolution, which was better than that of older adults. For classifications given during session 2, this led to a large effect of accuracy on z-scored confidence, χ^2^(1) = 137.1, *p* < .001, and a significant interaction between Accuracy and Age, χ^2^(1) = 12.9, *p* < .001. Both patterns held true for judgments made during session 3: a significant main effect of accuracy, χ^2^(1) = 164.2, *p* < .001, and a significant interaction between Accuracy and Age, χ^2^(1) = 25.0, *p* < .001. These results are consistent with findings showing that age-related metacognitive monitoring deficits in tasks involving recollective processes (for a review, see Dodson, 2017).

## Summary

Experiment 2 extended the findings from Experiment 1 in showing that detecting and recollecting changed activity features produced proactive facilitation in Day 2 recall, whereas detecting but not recollecting changes produced proactive interference. Conversely, detecting but not recollecting change increased Day 1 intrusions relative to detecting and recollecting change. These results confirm predictions of CCT that change recollection should moderate the effects of change detection by gating access to configural representations that preserve the temporal order of activity features. Experiment 2 also showed that both age groups could monitor the accuracy of the activity type classifications reasonably well, but older adults showed impaired resolution. These results are consistent with a larger literature showing age-related deficits in monitoring of recollective information.

### General Discussion

In the present experiments, we examined adult age differences in the detection and recollection of changes in everyday activities and memory for features of those activities. Our findings clearly showed that memory for activity features was better when changes were recollected than when they were not, and that this moderating effect of change recollection depended on change being detected first. Older adults showed a differential deficit in memory for changed activities that was partly due to their impaired detection and recollection of change. Older adults also showed poorer metacognitive monitoring in their classification of activity types, which likely resulted from the recollective demands of the task. The overall patterns of recall were consistent with predictions from CCT. We interpret these results further and discuss directions for theory development below.

### Detecting and Recollecting Everyday Changes

The primary goal of the present study was to investigate the consequences of memory updating following the detection of episodic changes. To accomplish this, we combined perspectives from the event perception and episodic memory literatures to create a controlled yet naturalistic procedure for examining these everyday cognitive phenomena. Our basic assumption was that effective parsing of perceptual inputs enables the comprehension of everyday activity features, which includes relationships between events and regularities in their occurrence. Experiment 1 showed that when change was recollected, Day 2 recall was enhanced and Day 1 intrusions were diminished. Experiment 2 extended these findings by showing that the effects of change recollection depended on change being detected first. In both experiments, proactive facilitation was obtained in Day 2 recall when change was recollected, whereas proactive interference was often obtained when change was not recollected. These results are consistent with accounts of episodic change processing in showing that comprehending changes in ongoing activity depends on successful recollection of configural traces.

The present results also showed that the beneficial effects of change recollection on recall were only obtained when participants could successfully recall Day 1 features. We interpret these results as showing that the quality of configural traces encoded during change detection depend on the accessibility of earlier memories. Such accessibility could vary across individuals and age groups depending on the perceived structure of events during encoding, which may determine the basis for retrieval during change detection. CCT assumes that configural traces are formed when Day 1 features are recollected and compared with current changed features. Following this, the representational structure of configural traces should differ depending on the accessibility of earlier features during change detection.

What determines the accessibility of features from previous episodes during comprehension of a new episode? Is it merely a matter of trace strength, or are there qualitatively different routes by which memory biases ongoing comprehension (Wahlheim, 2014)? Dual process theories posit that both consciously controlled (recollection) and automatic influences of memory contribute to retrieval across a variety of tasks (e.g., Jacoby, 1991; Yonelinas, 2002). Following this assumption, the retrieval of earlier events when detecting changes in current events may sometimes be accomplished through recollection or automatic influences. CCT proposes that change detection accomplished through successful recollection of Day 1 features should encode those features as part of a configural trace. Such a configural trace will have information about Day 1 features, Day 2 features, and the temporal relationship between them. The idea that recollection quality determines whether configural traces will reduce interference is consistent with the finding that hippocampal activity predicts the extent to which encoding of new memories is impaired by reactivating prior memories (e.g., Kuhl, Shah, DuBrow, & Wagner, 2010). Based on this, we suggest that recollection of event features during change detection will benefit order memory to the degree that Day 1 features are accessible when configural traces are retrieved. However, change detection accomplished through automatic retrieval should not confer these benefits because configural traces would lack the Day 1 features necessary to facilitate memory for Day 2 features. In the present experiments, one possibility is that when participants successfully recollected changes but were not able to retrieve Day 1 features, their change recollection was driven by automatic retrieval. (Of course, another path to correct change recollection responses is guessing, which surely also occurred.)

Consistent with the idea that proactive effects of memory may vary depending on the qualities of configural traces, the present experiments also showed that the magnitude of proactive interference effects when change was not recollected was smaller in Experiment 2 than Experiment 1. These findings are consistent with earlier studies showing that interference effects observed when change was not recollected increased when the accessibility of earlier memories was enhanced by repetitions (Wahlheim, 2014) and retrieval practice with feedback (Wahlheim, 2015). The present study showed that proactive interference effects were greater when each movie was shown in the same session (Experiment 1) rather than different sessions (Experiment 2). This difference in the temporal context associated with each movie across experiments may have resulted in events being more competitive in Experiment 1 than Experiment 2. However, fewer observations were available to compare recall when change was not recollected in Experiment 2, which indicates the need to verify these results using a direct within-experiment comparison. If this across-experiment difference is real, it parallels findings showing that context differentiation reduces interference (for reviews, see Abra, 1972; Smith & Vela, 2001). One consequence of this differentiation may be that features associated with distinct but related event models compete less when they are not retrieved as part of a configural trace.

In addition to differences in proactive effects of memory resulting from competition between encoding contexts, individuals may differ in their ability to encode Day 1 features as part of configural traces. An important and challenging proposal of CCT is that the relationship between encoding and retrieval is cyclical—at the same time that episodic retrieval is influencing ongoing comprehension, the mechanisms of comprehension are determining encoding for subsequent retrieval. Here, we examined individual age differences in change detection and their downstream effects on change recollection and recall performance. Older adults showed a differential deficit in Day 2 recall for changed activities. This deficit was associated with a poorer quality of trace integration, as shown by lower recall of Day 1 features for older than younger adults following change recollection. This age-related deficit may have reflected older adults detecting change on the basis of automatic influences more often than younger adults due to an age-related recollection deficit. Indeed, evidence for this was shown by Wahlheim (2014) in that older adults’ recall of earlier features following change detection was impaired, which led to poorer change recollection and greater interference in recall performance. Overall, these findings demonstrate how the negative effects of older adults’ impaired encoding of everyday activities (Kurby & Zacks, 2011; Zacks et al., 2006) can lead to downstream deficits in their detection and recollection of changes in those activities.

Older adults’ impoverished event representations may also have contributed to their impaired metacognitive monitoring of activity classifications. In Experiment 2, confidence judgments discriminated more poorly between correct and incorrect judgments for older than younger adults. This deficit in monitoring resolution may have partly resulted from lower-fidelity representations serving as less diagnostic bases for judgments. This is consistent with our earlier suggestion that older adults are less likely recollect prior events when viewing ongoing activities. This deficit is also noteworthy because earlier results (Wahlheim, 2014, Experiment 2) showed that confidence judgments were comparably sensitive to the effects of change recollection on recall performance in paired associate learning for younger and older adults. This was despite the fact that the retrieval task placed high demands on recollective processes. Together, these findings indicate that although older adults might show impaired metacognitive accuracy in their judgments regarding the relationships between events, they are aware of the effects of recollecting changes on recall. More generally, this discrepancy is similar to findings in the cognitive aging literature showing that even though older adults are more prone to false remembering (for a review, see Dodson, 2017), they are also fully aware of their memory deficits in many situations (e.g., Hertzog & Dunlosky, 2011). This highlights the need to illuminate the variables that moderate the relationship between age and metacognition of change processing.

### From Memory-for-Change to Change Comprehension

The present results replicate earlier findings that can be explained by the memory-for-change framework. But can the memory-for-change framework sufficiently explain the present results? No; the memory-for change framework is insufficient for explaining the present results because it does not account for the predictive processing involved in the perception of temporally structured events. As described in the Introduction, we propose that a merger between the memory-for-change framework and EST can provide a more comprehensive account (CCT) of how dynamic event changes are encoded and represented, along with the memorial consequences of recollecting those changes.

CCT assumes that the feature overlap between everyday events and prior event representations can cue involuntary or voluntary recollections of earlier events. We propose that such retrievals can serve to guide predictions of upcoming events, and that the perception of unexpected event features leads to prediction errors. These prediction errors should upregulate attention to unexpected features allowing for change detection and event model updating, consistent with predictions from EST (Zacks et al., 2007). Current event features should then be encoded with retrieved event features and associated cognitive operations into a configural trace, consistent with the memory-for-change framework (Jacoby et al., 2015; Wahlheim & Jacoby, 2013). Importantly, the formation of these traces requires that prior events be recollected while perceiving current events. The benefits of integrative encoding should be expressed when configural traces are accessed via change recollection. However, retrieval of recent related events during change detection should increase their accessibility, also making them a more robust source of interference when they are unopposed by change recollection.

Although the findings that we report here are consistent with CCT, the current experiments do not unequivocally demonstrate a role for memory-based predictive processing. However, research from the discourse comprehension literature provides support for the proposal that predictive processing plays a role in episodic change detection during everyday event perception. Findings from discourse comprehension research may generalize to event perception as both tasks may be governed by similar underlying processes. Evidence for this has been shown by situation models and event segmentation having similar effects on text and movie comprehension (Baggett, 1979; Lichtenstein & Brewer, 1980; Zacks, Speer, & Reynolds, 2009). Altmann and Kamide (1999) provide a clear example of the role of prediction in discourse comprehension. In their study, eye tracking was used to examine whether knowledge of semantic relationships would enable prediction of action plans when listening to sentences and viewing objects that corresponded to those sentences. They found that participants looked towards objects that were consistent with spoken verbs prior to those verbs being mentioned in the sentences. This was taken as evidence for memory-based anticipation of upcoming features (also see, Kamide, Altmann, & Haywood, 2003).

Research in the discourse comprehension literature might also inform potential age differences in the use of predictive processing in episodic change detection. For example, Federmeier, McLennan, DeOchoa, and Kutas (2002) used event related potentials to examine adult age differences in predictions of word features based on sentence context. They found that sentence context facilitated processing of within-category violations that were related to expected words for younger but not older adults. This was shown by N400 amplitudes being facilitated for within-category violations less effectively for older adults. These results suggest that older adults may be less able to construct representations that allow for the anticipation of upcoming features. Similar mechanisms may operate in event perception as older adults form less coherent event models that impair episodic retrieval and anticipation of activity features.

CCT asserts that episodic retrieval influences predictive processing, which in turn affects current encoding. A key challenge for future research is to separate the mechanisms of episodic retrieval and predictive processing in online comprehension. For example, one might directly assay memory retrieval using pattern-based functional MRI (Rissman & Wagner, 2012). In addition, to directly assay predictive processing, a promising approach is to measure anticipatory eye-movements prior to episodic changes. Individuals fixate to objects in naturalistic activities prior to when an actor makes contact with the object (e.g., Hayhoe, Shrivastava, Mruczek, & Pelz, 2003; Land & McLeod, 2000), and CCT predicts that this should be affected by retrieval from relevant previous episodes.

### Conclusion

The present study represents the first attempt to integrate perspectives from the event perception, episodic memory, and cognitive aging literatures to examine the detection and recollection of episodic changes in naturalistic activities. These findings open a new avenue for research that cuts across fundamental areas of inquiry in cognitive psychology and neuroscience. These results also demonstrate the viability of this approach for investigating the perception of and memory for changes in naturalistic activities in older and younger adults. Future directions involving a combination of cognitive and neuroscientific methods will allow us to further develop a comprehensive theoretical framework that attempts to explain how individuals update their memory to adapt to changes in the actions of others and how such updating might contribute to their own everyday activities.

### Context

This research grew from the intersection of research by Wahlheim and Jacoby on the memory-for-change framework with work by Zacks and colleagues on event representations in episodic memory. For us, a key theoretical insight was that the temporal dynamics of change detection are of major theoretical importance. CCT proposes that, during ongoing comprehension, recent episodic memory representations are retrieved, that the contents of these episodic memories inform one’s working models of the current activity, and that this informs predictions about what will happen in the near future. When such memory-guided predictions lead to errors, this induces a processing cascade that alters new memory encoding. Thus, retrieval and encoding are cyclically linked.

## Author Contributions

Both authors developed the study concept and contributed equally to the study design. C. N. Wahlheim programmed the experiments and supervised data collection and coding. J. M. Zacks performed analyses with input from C N. Wahlheim. C. N. Wahlheim drafted the manuscript, and both authors made revisions. Both authors approved the final version for submission.

